# Non-vectorial Integration of Intersectional Short-Pulse Stimulation Enables Enhanced Deep Brain Modulation and Effective Seizure Control

**DOI:** 10.1101/2025.01.02.631064

**Authors:** Tamás Földi, Miklos Szoboszlay, Zoltán Chadaide, Bence Radics, Bálint Horváth, Endre Vecsernyés, István Langó, Péter Ráfi, Andrea Pejin, Lívia Barcsai, Gábor Kozák, Nóra Forgó, Kristóf Furuglyás, Olivér Nagy, Anett J. Nagy, Tamás Laszlovszky, Zoltán Somogyvári, Magor L. Lőrincz, Orrin Devinsky, Antal Berényi

**Affiliations:** MTA-SZTE ‘Momentum’ Oscillatory Neuronal Networks Research Group, University of Szeged, Szeged, Hungary; Neunos ZRt, Szeged, 6725, Hungary; HCEMM-SZTE Magnetotherapeutics Research Group, University of Szeged; Szeged, 6720, Hungary; Department of Pathology, University of Szeged; Szeged, 6720, Hungary; Department of Neurology & Stroke, University of Tübingen, Tübingen, Baden-Württemberg, Germany; Hertie-Institute for Clinical Brain Research, Tübingen, Baden-Württemberg, Germany; Department of Physiology, Anatomy and Neuroscience, Faculty of Sciences University of Szeged; Szeged, 6726, Hungary; Department of Computational Sciences, HUN-REN Wigner Research Centre for Physics, Budapest, 1121, Hungary; Neuroscience Division, Cardiff University, Museum Avenue, Cardiff CF10 3AX, UK; Department of Neurology, NYU Langone Comprehensive Epilepsy Center, New York University, New York, NY 10016, USA; Neuroscience Institute, New York University; New York, NY 10016, USA

## Abstract

Transcranial electrical stimulation (TES) holds promise to treat neurological disorders, but its efficacy is limited by poor spatial focality and depth of penetration. Here, we examined the potential utility of Intersectional Short-Pulse (ISP) stimulation of deeper brain penetration. Using computational modeling and *in vivo* patch-clamp recordings in rats, we demonstrate that neurons integrate ISP-induced electric fields in a non-vectorial manner. This mechanism allows ISP to overcome some limits of conventional TES, achieving spatially limited stimulation across cortical and subcortical structures. In a rat model of temporal lobe epilepsy, closed-loop ISP stimulation significantly outperformed conventional TES in reducing seizure duration and severity. ISP reduced hippocampal seizure duration by 49% and 41% compared to sham stimulation and conventional TES and significantly reduced motor seizure severity. Our findings demonstrate that ISP stimulation can rapidly terminate hippocampal seizures, offering a potential new approach for non-invasive neuromodulation with applications across diverse neurologic and psychiatric disorders.

## Introduction

Transcranial electrical stimulation (TES) is a widely used non-invasive technique to modulate brain activity, a potential therapy for neurologic and psychiatric disorders (Begemann et al., 2020; Berényi et al., 2012; Brunoni et al., 2012, 2016; Chase et al., 2020; Fregni et al., 2006, 2021; Hawas et al., 2024; H.-H. Liu et al., 2019; Nitsche et al., 2008; Nitsche & Paulus, 2000; Priori, 2003; Stagg et al., 2018). By inducing weak electric fields in the brain, TES can alter neuronal transmembrane potentials and influence spike timing and network oscillations (Anastassiou et al., 2010; Antal et al., 2004; Nitsche, Liebetanz, et al., 2003; Nitsche, Nitsche et al., 2003; Nitsche & Paulus, 2000, 2001; Ozen et al., 2010; Priori et al., 1998; Purpura & McMurtry, 1965). However, the clinical efficacy of conventional TES is limited by poor spatial focality, limited penetration depth, and peripheral side effects at higher intensities (A. Liu et al., 2018; Poreisz et al., 2007; Vöröslakos et al., 2018).

Another technical challenge in TES is the "mirror effect," where simultaneous activation of surface electrodes induces opposing effects on brain activity under the cathode and anode (Creutzfeldt et al., 1962). This is particularly problematic when targeting spatially distributed oscillations generated by long-range neuronal networks, as in epileptic seizures (Berényi et al., 2012; Fusco et al., 2013; Gschwind & Seeck, 2016; San-Juan, 2021; San-Juan et al., 2018; Zoghi & Jaberzadeh, 2021). Electrode configurations to address this issue fail to eliminate the mirror effect or overcome the trade-off between focality and depth of stimulation (Datta et al., 2009).

Innovations in transcranial electrical stimulation (TES) have increasingly focused on enhancing the spatial precision and depth targeting of applied electric fields. Spatial summation of multiple current sources is a promising strategy, exemplified by methods like high-definition tDCS (HD-tDCS) and Temporal Interference (TI) stimulation. HD-tDCS employs arrays of small electrodes to confine the electric field to targeted cortical areas, improving focality and minimizing off-target effects (Datta et al., 2009; Dmochowski et al., 2011; Hogeveen et al., 2016; Tedla et al., 2023; Villamar et al., 2013). TI stimulation, introduced by Grossman et al., generates low-frequency electric fields at specific brain regions through interference patterns of high-frequency currents, enabling to constrain the interference part of the stimulation (i.e. the beat frequency) to deeper brain regions (Demchenko et al., 2024; Grossman et al., 2017, 2018; Luff et al., 2024).

Both HD-tDCS and TI represent substantial advancements in spatially directing the effects of brain stimulation. However, achieving higher stimulation intensities for enhanced deep-brain modulation while minimizing peripheral and off-target side effects remains an ongoing challenge.

Maxwell’s theory of electromagnetism and the principle of superposition hold that the electric field distribution within a conductive volume (e.g., the head) is determined by the net current flow, whether this current originates from multiple independent sources or a single common source connected to multiple electrodes. For TES, when multiple independent current sources are attached to the scalp, the electric field distribution and strength within the head is identical to what would be produced by a single source connected to the same electrode montage if the net current input at each electrode location is the same in both scenarios. Thus, using multiple independent current sources instead of a single source with multiple electrodes does not improve the electric field’s focus or shape in the brain (Figure 1; Supplementary Figure 1;). Any technique that enhances spatial focality must rely on principles or methods beyond merely increasing the number of independent current sources.

**Figure 1.**
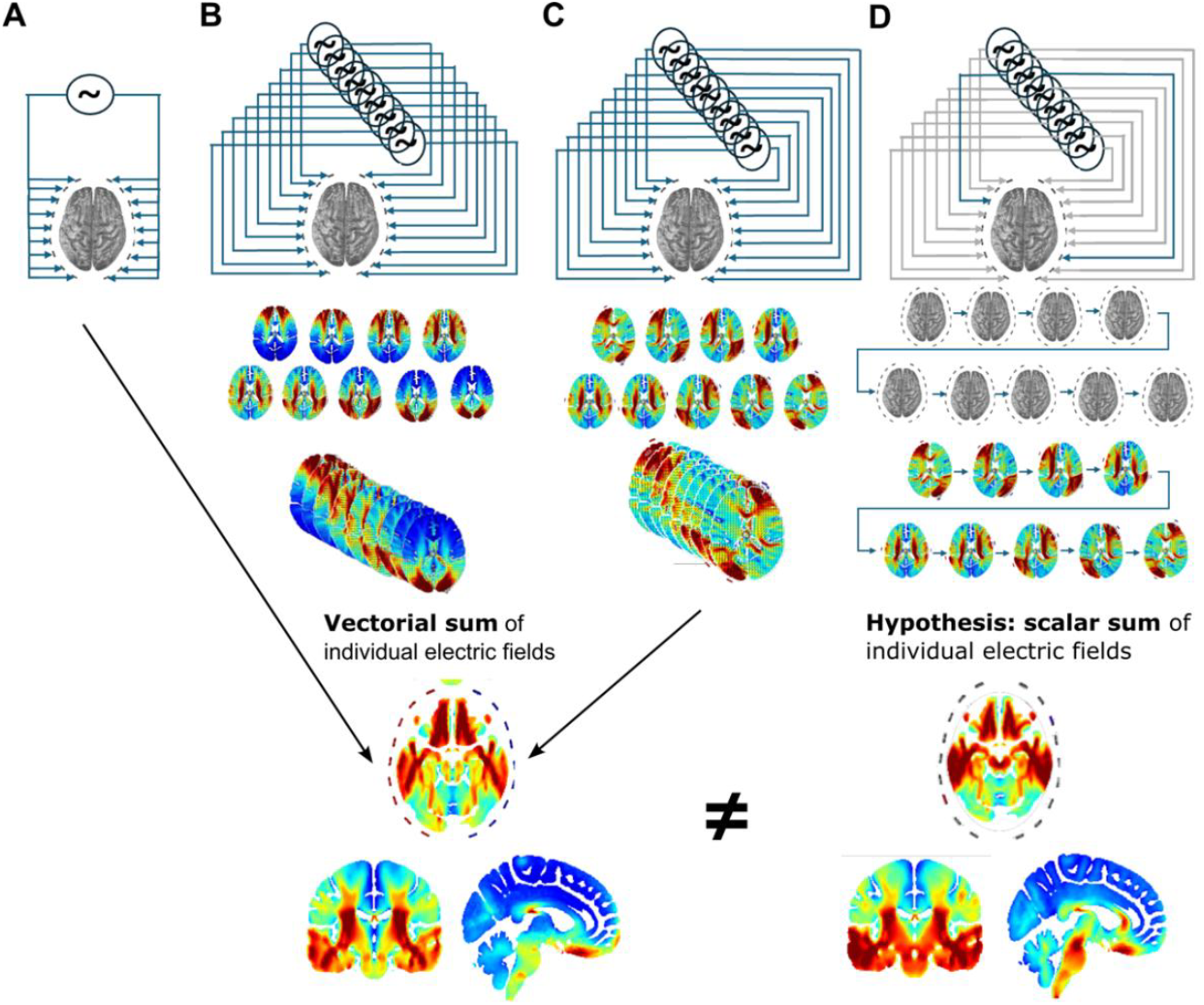
Equivalence of electric field distributions in transcranial electrical stimulation (TES) under different source configurations. This figure illustrates the principle that the electric field distribution in the brain is determined by net current flow, regardless of the source configuration. Three scenarios are presented: **A**): A single common source driving all electrodes simultaneously, similar to HD-tDCS. **B**): Multiple independent sources applied simultaneously with non-crossing current paths, similar to TI stimulation. **C**): Multiple independent sources applied simultaneously with crossing current paths. The top row shows schematic representations of the electrode placements and current paths of the individual sources for each scenario. The bottom row displays the simulated electric field distributions in axial, coronal and sagittal views. All three scenarios produce identical electric field distributions within the brain since the principle of superposition dictates a geometric (i.e. vectorial) net effect as the sum of independent sources if the total current input at each electrode is equivalent across scenarios. In contrast, if the independent sources are sequentially but not concurrently activated (**D**), the electric fields remain discreet, and integration occurs intracellularly as charge accumulation. Due to the non-linearity of the transmembrane charge injection, we hypothesize that this arrangement results in a different neuronal ‘readout’, which more closely resembles the algebraic (i.e. scalar) sum of individual electric fields.

An alternative to the spatial summation is to exploit the time-integrative properties of the neuronal membrane. When multiple source effects are time-integrated by neurons by applying independent sources in rapid succession rather than simultaneously (Figure 1D), each discreet extracellular electric field injects current across the neuronal membrane. Since transfer across the cell membrane is non-linear, neurons can integrate individual charge injections induced by these rapidly successive, quasi-independent source effects.

The principle of intracellular integration has several implications. The extracellular space is largely an ohmic conductor, causing the extracellular fields to build up and disappear instantaneously. By applying electric fields in rapid succession rather than simultaneously, each field induces its discrete charge injection before the next field is applied. The cell membrane’s capacitive properties enable it to accumulate charge over time, even from fields that would normally cancel in the extracellular space due to their vectorial summation by superposition. Further, the non-linear characteristics of the membrane’s transfer function may amplify or modulate the integration of charge injections in ways not predictable from linear superposition.

Using the time-integration principle, Intersectional Short-Pulse (ISP) stimulation (Vöröslakos et al., 2018) was introduced to increase intracerebral field strength while reducing the adverse peripheral effects. The ISP approach can circumvent the superposition principle as applied to electric fields in the extracellular space. This enables non-invasive brain stimulation with more precise and effective neuromodulation in research and clinical applications.

To test the predictions of the ISP hypothesis, we combined computational modeling, cadaver measurements, *in vivo* electrophysiology, and a rodent model of temporal lobe epilepsy. We elucidate the mechanism of ISP stimulation by membrane integration in *in vivo* intracellular recordings and demonstrate its superior efficacy in controlling epileptic brain activity.

## Results

### Validation of ROAST to high-frequency ISP stimulation

ROAST (Realistic volumetric-approach to simulate transcranial electric stimulation) is an established finite element modeling software that models the intracerebral electric potentials and potential gradients (i.e. electric fields) during quasi-static tDCS-like transcranial electrical stimulation (Huang et al., 2016, 2019). While the primarily ohmic conductance of the brain’s extracellular space is demonstrated in the sub-MHz frequency range (Vöröslakos et al., 2018), ROAST has not been validated for ISP with instantaneous electric fields with rapidly changing pulsed stimuli.

ISP stimulation sequences were studied with brief, discrete tDCS-like stimuli and varying electrode arrangements applied to two human cadaver heads through four 8-channel subgaleally implanted electrode strips (32 contacts). Intracerebral potentials were measured by eight 8-channel stereoEEG (sEEG) electrodes implanted geometrically regularly on cadaver brains through small, isolated burr-holes, maximally maintaining scalp and skull integrity (Figure 2A).

**Figure 2:**
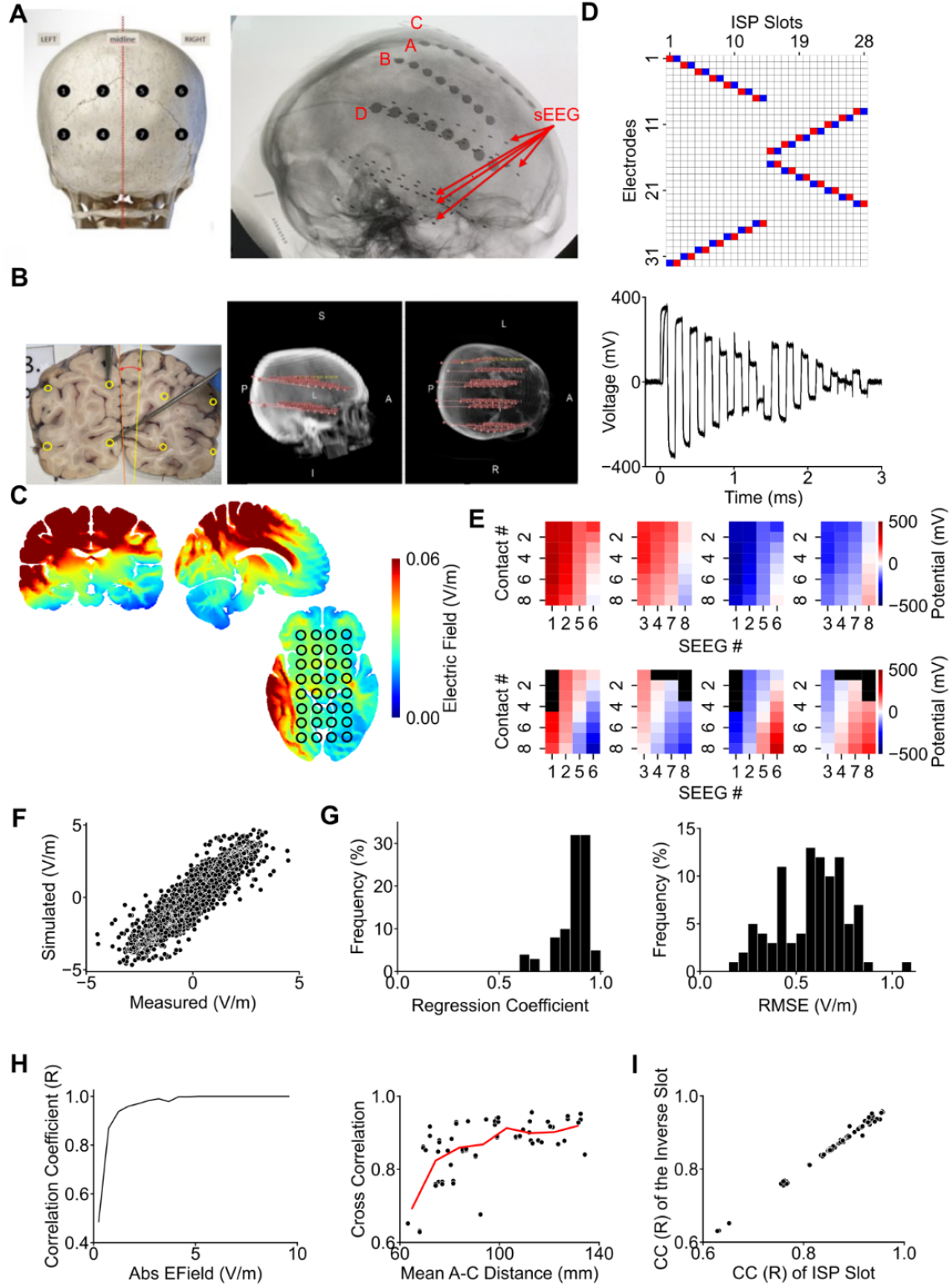
Validation of ROAST finite element modeling for ISP stimulation in human cadavers. **(A)** Experimental arrangement of simultaneous intracerebral sEEG recording and transcranial ISP stimulation in human cadavers. Four eight-contact stimulation strips, labeled C, A, B, and D from left to right, are implanted subgaleally, parallel to the midline. Contact points of each electrode strip were numbered from one to eight, from the front to the back. **(B)** An example brain slice of the post-experimental tissue processing, showing the stained electrode tracks used for post-hoc reconstruction of the sEEG recording electrode locations. 3D image shows the 3D reconstruction of the verified recording sites on the MR image of the head. **(C)** An example simulated effect of one ISP slot is shown as heatmaps on the cross-sections of the three main plains of the brain. Marks on the heatmaps denote these locations, from where the simulated potential levels were extracted. **(D)** The plot shows the individual voltage readouts of one intracerebral sEEG electrode contact, superimposed on each other, during the "Central" stimulation sequence. Note that the different electrode assignments within the sequence induced distinct potential readouts throughout the stimulation sequence. The electrode assignments to each ISP stimulus pattern (‘slot’) are shown above the recording. To ensure charge balancing, every other electrode assignment is identical to the preceding assignment with the polarity flipped. Blue: cathode, Red: anode, White: disconnected. **(E)** Measured (top row) and simulated (bottom row) potential maps generated by the simultaneous readouts of all sEEG electrodes during four example individual ISP pulses. Black: excluded electrodes due to unrecoverable spatial location in the brain. Note the similarity of the voltage gradient patterns of the corresponding measured and simulated pairs despite the shifted reference (i.e. zero) levels of the measured maps. **(F)** Correlation of the simulated and the measured voltage gradients of all possible sEEG electrode pairs in an example ISP Stimulation pattern. **(G)** Distribution of regression coefficients (left panel) and RMSEs (right panel) of the correlations of the measured and simulated electrical field gradients across all ISP stimulation patterns. **(H)** Correlation of measured and simulated electric field gradients as a function of gradient magnitude (left) and average anode-cathode distance (right). **(I)** Similarity of the correlation coefficients of ISP slots with identical stimulation parameters but inverted electrode polarities.

A custom transcranial neurostimulator (Neuroclinical, Neunos Ltd, Szeged, Hungary) delivered charge-balanced sequences of ISP stimuli. Individual ISP stimulus pulses were 100 μs long, typically 24-30 stimuli were delivered in a sequence. All ISP sequences were repeated ≥100 times, then averaged to improve the signal-to-noise ratio (SNR) of intracerebral electric field recordings. Four ISP sequences were designed to target frontal, parietal, or occipital lobes or midbrain.

The exact subgaleal contact site locations used for stimulation were mapped on the international 10/10 system. The post-experimental processing of the brains reconstructed the dye-covered sEEG electrode tracks and contact site locations. Locations were transformed into recording site voxel coordinates (Figure 2B).

The cadaver head MRIs were used to model the induced intra-cadaver brain potential distributions for each ISP stimulation slot (i.e. for each electrode assignment and stimulation arrangement) by a modified version of the ROAST modeling environment, allowing for subgaleal stimulation electrode placement. This iterative modeling approach led to a set of instantaneous potential distribution maps, one for each brief stimulus pulse in the stimulation sequence. The predicted potential levels for each SEEG contact site were extracted from the FEM models (Figure 2C).

The induced intracerebral electric fields were recorded at 64 or 80 locations by a high-speed DC-coupled electrophysiologic amplifier (KJE-1001, Amplipex, Szeged, Hungary) at 1.28 MS/s sampling rate (Figure 2D). Baseline shifts of the intracerebral recordings due to any direct current biopotentials were removed *post hoc*. The potential traces were discretized by reading out the potential value of each contact point at the late phases of each ISP slot to avoid the effects of switching transients.

The discretized values of all sEEG contact points were used to calculate the measured potential gradient maps for each ISP stimulation pattern, and the potential values of the corresponding voxels of the simulations were extracted (Figure 2E). Some electrode locations were unidentifiable by the *post hoc* histological analysis; these electrode potential values were excluded from the analyses (N = 9 of the 64 electrodes in two cadavers). The reconstructed contact site locations informed the inter-electrode distances to calculate the electric fields from the contact site potential values during the processing of the simulation and experimental data and predicted electric fields.

The correlation of simulated and measured inter-electrode voltage gradient pairs (2016 to 3160 data points per ISP pattern, 94 ISP total patterns) was calculated (RMSE; Figure 2F). The modeled and measured voltage gradients showed a high correlation (median R = 0.8791, IQR = 0.079, p < 0.001; median RMSE = 0.6048, IQR= 0.2613; N = 94 for both comparisons), also validating the ROAST model for transient electric fields (Figure 2G). The detailed analysis of the data points with lower correlation revealed a larger average distance between anodes and cathodes, and larger induced electric field gradients improved model predictions, while stimulation between electrodes in close proximity (e.g. between neighboring electrodes on the same electrode strip) decreased model prediction (Figure 2H), likely reflecting stronger local tissue shunting and higher sensitivity of the model on the exact tissue segmentation and MRI resolution. The modeling framework’s reliability is also supported by the consistent correlation coefficients for ISP slot pairs where electrode polarities were swapped, but the electrode assignments were identical (Figure 2I, Correlation of correlation coefficients: R= 0.9973, p < 0.00; Correlation of RMSEs: R= 0.8874, p < 0.00; N = 46 ISP slot pairs, in both comparisons). In conclusion, these measurements confirm the validity of ROAST as a modeling environment for the rapidly switching pulses of ISP. This makes it an appropriate tool for modeling the local electric field vector sequences of ISP, facilitating further modeling of neuronal readouts.

### Modeling the neuronal response to ISP stimulation

To investigate how individual neurons integrate ISP stimulation, we simulated neuronal responses of ‘humanized’ model neurons with realistic morphologies (Figure 3A, (Aberra et al., 2018)) by exposing them to simulated electric fields generated by ISP and conventional TES stimulations. The rheobase of the neurons (i.e., minimum voltage gradient capable of inducing an action potential (AP)) was tested as a function of the resting transmembrane potential and neuronal orientation. We found a steep decrease in excitability when neurons were hyperpolarized below -86 mV, a close to a linear relationship between -63 and -86 mV, and sporadic spontaneous APs above -62 mV (Figure 3B). Neuronal excitability depended on the orientation of the neurons, resulting in an ∼3.5-fold difference (median = 3.57, IQR = 7.93, N = 25) in the rheobase between the maximal and minimal excitable orientations (example neuron on Figure 3C: 32 V/m at θ = 120° and φ = 60° vs. 10 V/m at θ = 270° and φ = 90°, 0.9 ms-long square wave stimulus, L2/3 pyramidal neuron, resting membrane potential -73 mV). While the subthreshold stimulus often induced somatic hyperpolarization, increasing the stimulation intensity consistently induced spiking (Figure 3C). In line with previous modeling studies, our *in silico* results showed that extracellular stimulation elicits heterogeneous V_m_ changes in a compartment-specific manner. APs can be initiated in distal subcellular regions while the soma is hyperpolarized so that axonal/dendritic action potentials spread to the soma and overcome the initial local hyperpolarization, resulting in somatic firing (Rattay et al., 2017).

**Figure 3:**
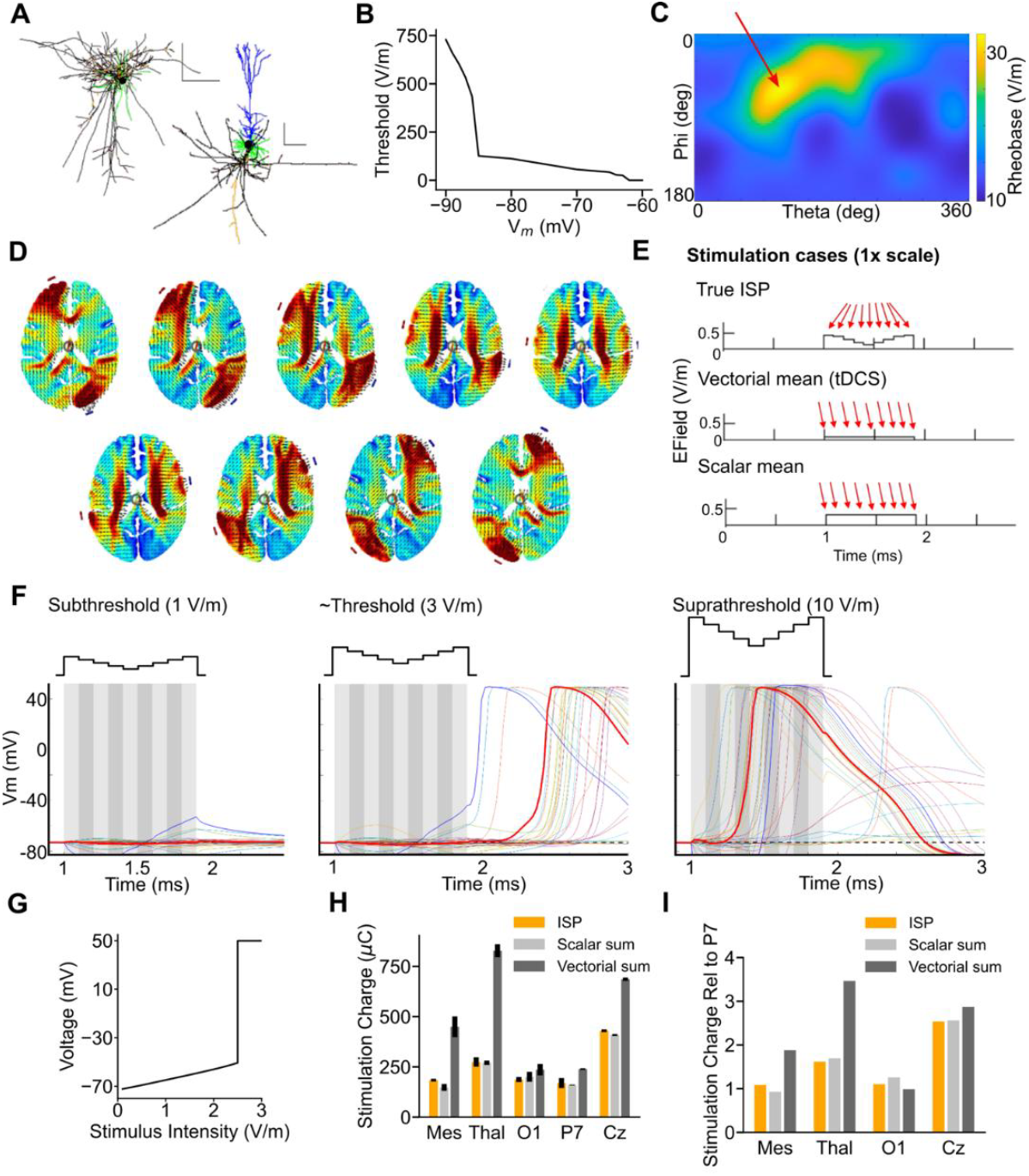
Simulation of ISP and diffuse transcranial electrical stimulation effects on single neurons in the human brain. **(A)** Neuron model morphologies (scale bars: 50 μm). **(B)** The stimulus threshold to induce an AP with ISP stimulation strongly depends on the resting membrane potential of the neuron, particularly at strongly hyperpolarized states. **(C)** Altering the sources’ direction (i.e. anodal vs. cathodal) caused only a three-fold change in the required current intensity to evoke an action potential. Arrow denotes the least excitable orientation of the example neuron. **(D)** Modeled instantaneous electric field maps of consecutive slots for a 9-slot rotating ISP sequence. **(E)** Magnitudes and directions of consecutive 100 µs-long electric fields were used as stimuli in the three stimulation scenarios. Figure shows the gradually changing directions and magnitude of the true ISP stimulation (top) and the constant vectors applied as the vectorial (middle) and scalar mean (bottom) equivalents. **(F)** An example of the neuronal responses for a single 0.9 ms-long sweep of the true ISP stimulation sequence. Thick red lines denote the somatic transmembrane potentials. Note that while an electric field pattern of 1 V/m average intensity failed to induce an action potential, the 3 V/m fields evoked an action potential at the soma 1.4 ms after stimulus offset, while a 10 V/m field evoked an instantaneous AP at the soma. **(G)** The highest local depolarization at any of the segments of the model neuron as a function of stimulation intensity. Note the linear relationship at subthreshold intensities. **(H)** The stimulus intensity threshold for ISP entrainment (orange) is similar near the electrodes (auditory cortex, AuC) and at the targeted deep crossing points (left and right hippocampi). The integrated ISP effect is best approximated with the scalar sum of individual ISP pulses (light grey) than their vectorial sum (dark grey), which requires substantially higher intensity to entrain deep structures than at sites proximal to the stimulating electrodes. **(I)** Same as (G), relative to surface-proximal structures (P7).

We next modeled the electric field maps generated by each ISP slot of a rotating ISP stimulation pattern, consisting of nine different electrode configurations (Figure 3D). The field vectors were extracted at five arbitrarily chosen anatomical locations (i.e. Mesencephalon, thalamus, visual cortex (at O1), parietal cortex (at P7), and at the Cz location). The nine vectors extracted at a location were used to construct the 9 ×100 µs long stimulation pattern applied in the NEURON simulations. We also constructed two tDCS equivalents of the ISP, with the magnitude of the vectorial and scalar mean of individual ISP field vectors and with their average direction (Figure 3E). The model neurons were aligned to their least excitable orientation to the mean direction of the electric field vectors, and their resting membrane potential was at -73 mV.

We determined the rheobase of the model neurons for all three stimulation types (i.e. ISP and its vectorial and scalar mean tDCSs) by measuring their membrane potential (V_m_) readouts at incrementally increased stimulus intensity (Figure 3F, G).

During ISP stimulation, the spike threshold intensity was similar for neurons near the electrodes and those at targeted subcortical areas (229 ± 54.11 µC/ms vs 262 ± 130.87 µC/ms, N = 4 vs 6, for superficial and deep targets, respectively, Fig. 3H). In contrast, the application of the vectorial sum of the ISP-generated fields, resembling their simultaneous application as conventional TES, required 65 ± 5 % higher stimulation intensities to entrain deep structures compared to cortical areas proximal to the electrodes (387 ± 231.99 µC/ms vs 639 ± 221.34 µC/ms, N = 4 vs 6, for superficial vs deep targets, respectively, Fig. 3I). This finding supports the hypothesis that ISP can achieve more uniform stimulation across different brain depths.

Further, our model indicated that the neuronal response to ISP stimulation correlated better with the scalar sum of the individual ISP fields rather than their vectorial sum (Fig. 3H and 3I, R= 0.98, p < 0.001 for ISP vs scalar sum, and R= 0.71, p = 0.023 for ISP vs vectorial sum, N = 2 × 10 in both comparisons, Pearson’s linear correlation), providing computational support that neurons integrate ISP-induced fields in a non-vectorial manner.

To evaluate the spatial effect of ISP stimulation, we incorporated a universal rat head model (Garcia-Gonzalez et al., 2018) into the ROAST toolbox (Huang et al., 2016, 2019). We also simulated the volumetric electric field distribution of our stimulation protocols in rats and channeled the electric field readouts to the same neuronal models as above. These results were qualitatively comparable to the simulations performed on the human brain. They supported the non-vectorial summation of the ISP stimulation effects and better entrainment of deep structures than conventional TES (Supplementary Figure 2.).

### *In vivo* validation of non-vectorial field integration

To validate our computational findings on the non-vectorial integration of the momentary effects of the subsequent electric fields, we performed *in vivo* whole-cell patch-clamp recordings in layer 2/3 pyramidal neurons of the rat somatosensory cortex during ISP stimulation (Fig. 4A). We recorded intra- and extracellular potentials to calculate transmembrane potential changes in two cells, while concurrently applying ISP stimulation (Fig. 4B). The ISP stimulation sequence was designed to be symmetric and charge-balanced, i.e. alternating pulses and their evoked electric fields had the same magnitude but inverse directions (Fig. 4D). For vectorial summation, the stimulus pair effects would cancel each other, and would not change the transmembrane potential.

**Figure 4.**
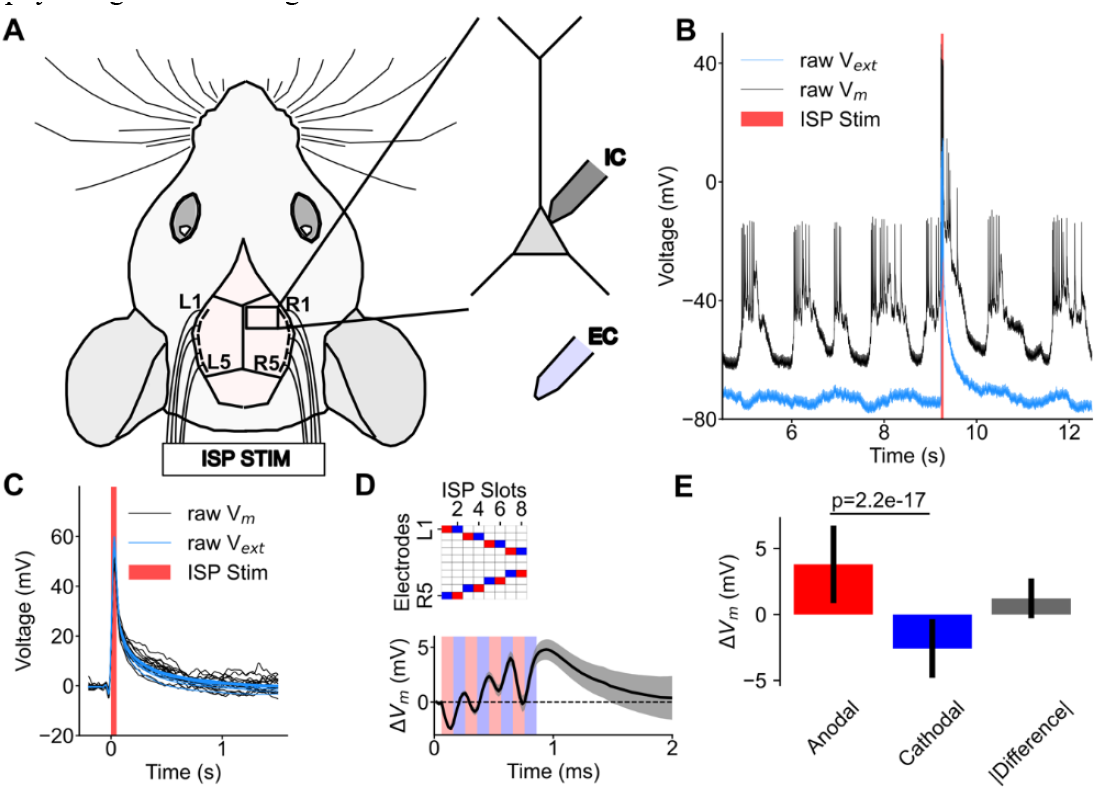
In vivo transmembrane potential recordings in rats during symmetric ISP stimulation. **(A)** Experimental arrangement using five pairs (L1-L5 and R1-R5) of transcranial stimulation electrodes and an intracellular and subsequent extracellular recording. **(B)** Scheme of the two-step measurement to eliminate the stimulation artifact. **(C)** Intra-(IC; black) and extracellularly (EC; blue) recorded voltage changes of the recorded cells triggered by ISP stimulation (red shaded area). **(D)** Transmembrane potential changes induced by the individual ISP pulses. The table above denotes the stimulation sequence and electrode assignment. Note the perfectly symmetric and charge-balanced stimulation pattern. **(E)** The average V_m_ changes induced by the anodic and their mirrored cathodic pulses and their difference (right panel). Note the non-zero-sum of symmetric pairs supporting a non-linear and non-vectorial summation.

To eliminate the seconds-long ISP-induced electrical artifact from the electrophysiological recordings and evaluate its evoked net effect, the patch-clamp recordings were followed by extracellular recordings from the vicinity of the recorded neuron (Figure 4A, B). In both configurations, ISP sequences were repeated 17-20 times and averaged (Figure 4C). The mean extracellular voltage waveform was subtracted from the mean intracellular trace, eliminating the common mode stimulation artifact and resulting in the stimulation-induced net transmembrane potential change (Figure 4D).

In both recorded cells, 0.8 ms of ISP stimulation at 100 μA triggered a net depolarizing effect, confirming that the symmetric stimulus pulse pairs induce non-symmetric changes in the transmembrane potential (Fig. 4D). Our analysis at a higher temporal resolution revealed that the anodic stimulus pulses elicited a stronger membrane potential change than its cathodic counterpart, resulting in a cumulative net depolarization (3.79 ± 2.94 mV for anodal and -2.58 ± 2.21 for cathodal stimulation, p < 0.001, paired t-test between the absolute values, N=2×144 value pairs, Fig. 4D and E).

We applied the same modeling pipeline described previously to mechanistically analyze the neuronal responses to symmetric ISP stimulation. The instantaneous electric fields induced by the individual electric pulses were simulated using ROAST (Supplementary Fig 3A), and subsequently incorporated into the NEURON model. The results revealed similar asymmetric transmembrane potential dynamics in response to symmetric stimulation, consistent with our *in vivo* measurements. A detailed analysis of the individual neuron segments showed that specific compartments were more sensitive to stimulation than the soma (Supplementary Fig. 3B). In many cases, stimulation near the cell’s rheobase intensity triggered action potentials in these distal segments rather than at the soma or axon hillock (Supplementary Fig. 3C).

These *in vivo* findings corroborate our computational results, supporting that integration of ISP pulse effects results from the cell membrane’s capacitive properties rather than extracellular integration of electric fields. This integration is non-vectorial.

### Spatially focused ISP stimulation

Previous findings demonstrated that precisely timed electrical stimulation (Figure 5A) can efficiently interfere with neuronal oscillations and terminate thalamocortical seizures (Berényi et al., 2012; Kozák & Berényi, 2017). However, closed-loop diffuse TES was less effective in terminating tonic-clonic seizures in a bilateral TLE model. Our simulations support the hypothesis that ISP stimulation can generate electric fields with comparable strength and direction in both hippocampi, unfeasible with conventional TES (Figure 5B). ISP also allowed for the generation of local electric gradient vectors with multiple orientations in each hippocampus, potentially amplifying local effects due to the complexity of neuronal morphology.

**Figure 5:**
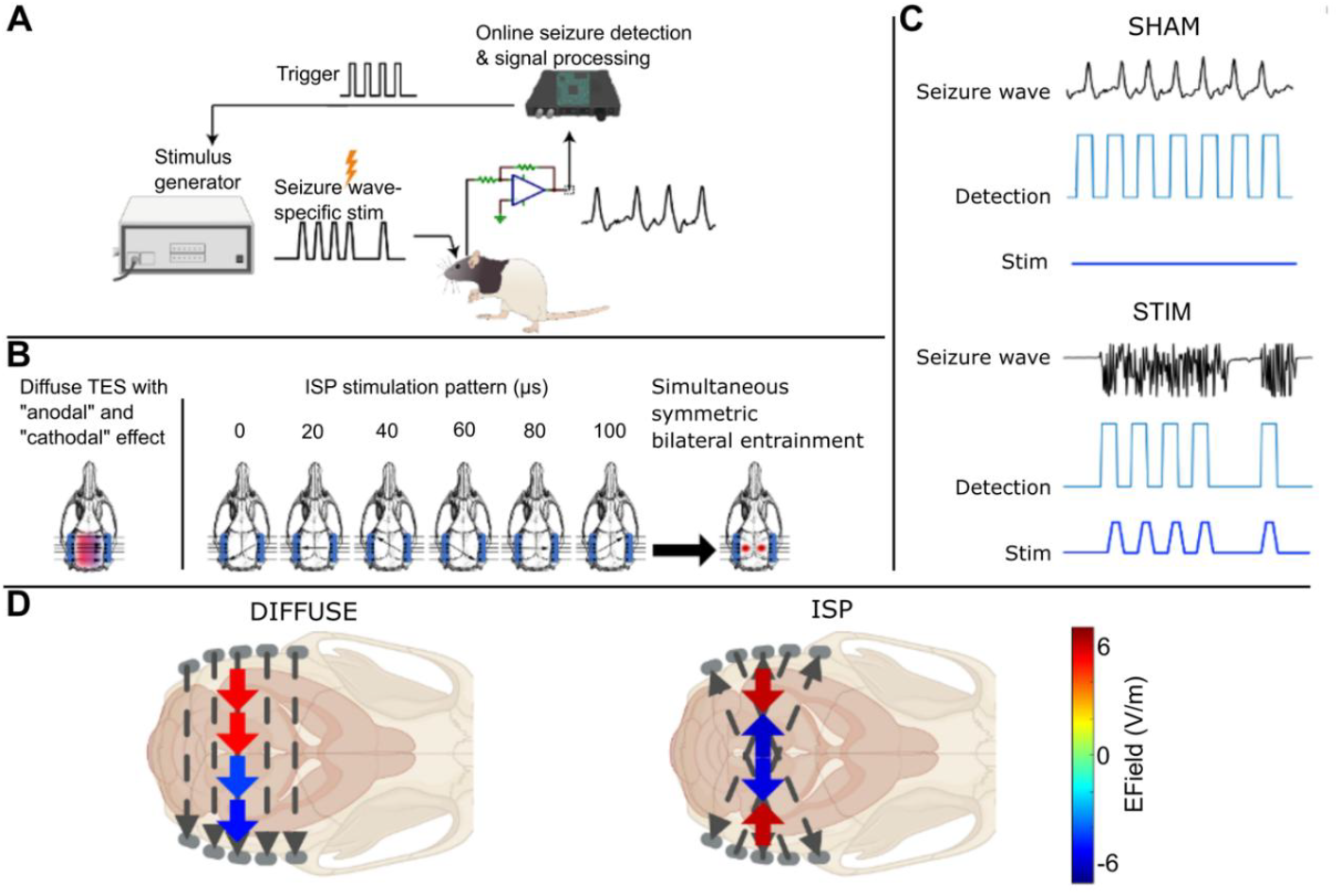
Closed-loop intervention and ISP stimulation sequence. **(A)** The concept of closed-loop seizure intervention through seizure rhythm-driven stimulation. Individual seizure waves were detected in real-time and each detection triggered transcranial stimulation. **(B)** Stimulation protocol of diffuse TES (left) compared to ISP (right). In the latter case, three electrode pairs were set to target the ISP pulses on either the left or the right hippocampus. Each electrode pair was pulsed for 20 μs, and the pulses cycled through the three pairs; this sequence was repeated to alternate between stimulating the right or left hemisphere, respectively. **(C)** A representative HPC seizure, its detection in the absence and presence of transcranial stimulation, upper and lower panels, respectively. **(D)** Measured average intracerebral gradients generated by diffuse stimulation (left) and ISP stimulation (right). Arrows denote the direction of the generated average electrical fields between the probe electrodes, while their colors denote their magnitude (5.29, 5.367, -5.18, -4,47 V/m for diffuse, and 6.011, -6.11, -6.033, 5.71 V/m for ISP stimulation) during one stimulation cycle shown in the horizontal plane.

To test if ISP is effective in controlling network oscillations, we used hippocampus-kindled rats as a model refractory temporal lobe epilepsy (TLE model; Gelinas et al., 2016) and attempted to terminate stimulation-induced bilateral TLE seizures by inducing a closed-loop symmetric bilateral entrainment. Rats (*n*=8) were kindled by daily electrical stimulation of the hippocampal commissure. The stimulation immediately provoked 10–25 Hz after-discharges bilaterally in the hippocampus. The kindled seizures were secondarily generalized (Racine’s score (RS) 5 motor seizures (8/8 rats). As positioning stimulating electrodes based on novel rat skull models (Goerzen et al., 2020) can increase effective spatial targeting, we incorporated a universal rat head model (Garcia-Gonzalez et al., 2018) in the ROAST toolbox (Huang et al., 2016, 2019; Nielsen et al., 2018) to model the volumetric electric field distribution of our stimulation (Figure 5A).

We used two closed-loop transcranial stimulation paradigms to terminate seizures: ISP or convectional (diffuse) TES. TES protocols were applied between the left and right temporal screw electrodes, placed onto the frontal and temporal bone (Fig. 5B, left). During ISP stimulation, we used six rotating dipoles to focus both hippocampi (Fig. 5B, right). The stimulus pulse intensities were identical in both cases and were triggered by ictal LFP deflections in the hippocampus, resulting in continuous electrical stimulation throughout the ictal period (*Figure 5A and 5C*). The signals were analyzed online to detect each LFP deflection in the HPC using a custom-made seizure detection algorithm built on an existing routine (Kozák & Berényi, 2017). We applied traditional TES using trapezoidal waveforms following the detection of paroxysmal LFP events. The induced hippocampal seizures were detected online, and each HPC LFP deflection triggered a single-pulse stimulation delivered either as ISP or as conventional TES (+8 V trapezoid, 80 ms duration) (*Figure 5A and 5C*).

Compared to diffuse TES, ISP stimulation showed more heterogeneous electric fields throughout the brain. While technical limitations of the experiment only allowed for a limited spatial resolution along one axis, we found that while the field directions were consistent and pointing from one side to the other in the case of the diffuse stimulation, for the ISP stimulation the field vectors were pointing towards the target zones (Fig. 5D). This critical refinement in spatial targeting improved on-target and reduced off-target neuromodulation effects.

### Efficacy of closed-loop ISP stimulation in terminating temporal lobe seizures

To test the efficacy of ISP in controlling pathological brain activity, we employed a hippocampal kindling model of TLE using male Long-Evans rats kindled by daily electrical hippocampal commissure stimulation until they consistently exhibited stage 5 RS seizures. We compared the effects of closed-loop ISP stimulation to conventional TES and sham stimulation on seizure duration and severity.

Closed-loop ISP stimulation significantly reduced normalized hippocampal seizure duration compared to sham (50.92±0.79 % vs 99±0.39 %, p<0.0001) and conventional TES (50.92±0.79 % vs 86.71±0.58 %, p<0.0001) (Fig. 6B1). Similar results were observed for cortical seizure duration (Fig. 6B2).

**Figure 6:**
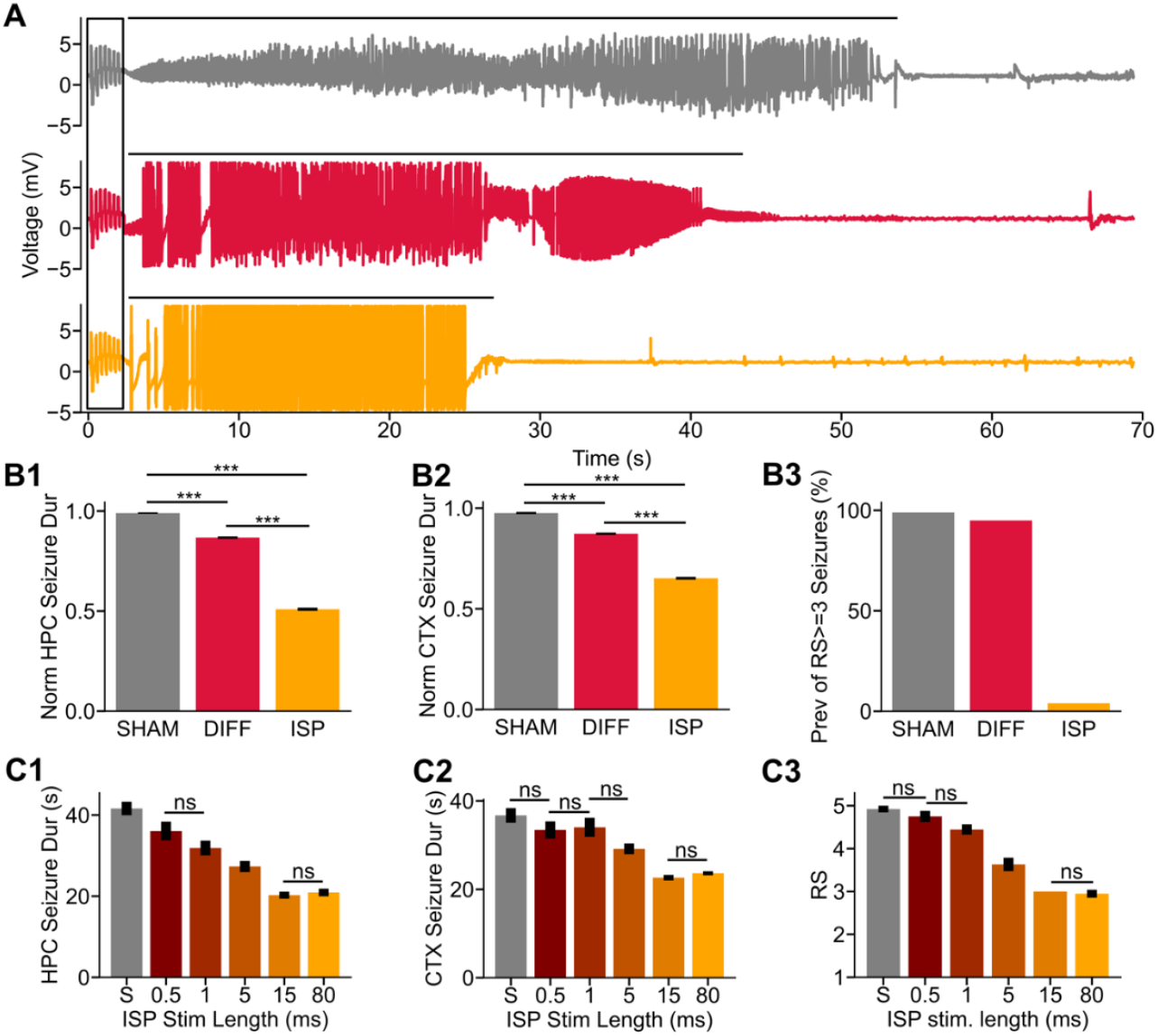
Anti-epileptic effects of ISP stimulation. **(A)** Representative LFP traces of HPC seizures induced by kindling stimulation with or without closed-loop transcranial stimulation. The black bracket denotes the kindling epochs, while black lines represent detected seizures. **(B)** Normalized (to the longest seizure of the day of recording) HPC seizure duration during SHAM (grey), DIFFUSE (red), and ISP (orange) stimulation (B1) (p< 0.0001, Wilcoxon signed-rank test; n=210 in 8 animals). Normalized CTX seizure duration during SHAM (grey), DIFFUSE (red), and ISP (orange) stimulation (B2) (p< 0.0001, Wilcoxon signed-rank test; n=210in 8 animals). Percentage of >=3 RS seizures during SHAM (grey), DIFFUSE (red), and ISP (orange) stimulation (B3) (99%, 95%, and 4%). **(C)** HPC seizure duration during different ISP stimulations of various durations (C1) (post-hoc Nemenyi test; n=24 in 2 animals). CTX seizure duration during different lengths of ISP stimulation (C2) (post-hoc Nemenyi test; n=24 in 2 animals). Racine’s score during different lengths of ISP stimulation (C3) (post-hoc Nemenyi test; n=24 in 2 animals).

ISP stimulation substantially reduced the severity (RS) of seizures (ISP: 2.5283±0.5628, diffuse TES: 4.3352±0.4734, sham: 4.8793±0.3267, all pairwise comparisons p<0.0001) (Fig. 6B3). This marked reduction in seizure severity demonstrates the superior efficacy of ISP in reducing seizure intensity compared to traditional TES.

We optimized the ISP stimulation protocol by systematically varying pulse durations. Decreasing stimulation duration from 80 to 15 ms did not affect efficacy, while below 15 ms effectiveness decreased (Fig. 6C). This optimization allowed for minimizing stimulation duration to achieve the same effect but reducing unwanted increased stimulation of individual sites, thus decreasing side effects and improving tolerability.

Together, these results demonstrate that closed-loop ISP stimulation can rapidly and effectively decrease the duration and severity of temporal lobe seizures more effectively than conventional TES, further underlying its potential as a tool for deep brain stimulation.

## Discussion

Our data demonstrate the superior efficacy of Intersectional Short-Pulse (ISP) stimulation in terminating epileptic seizures and elucidate its mechanistic differences from conventional transcranial electrical stimulation (TES) in entraining neuronal activity. We demonstrate that ISP has superior spatial focusing and efficacy in modulating neural activity versus conventional TES, particularly for terminating temporal lobe seizures. The key mechanism underlying ISP’s effectiveness is the non-vectorial integration of induced electric fields by neurons, allowing for deeper and more uniform stimulation while minimizing peripheral side effects.

### Advancements in Transcranial Electrical Stimulation

The limitations of conventional transcranial electrical stimulation (TES), including poor spatial focality, shallow depth penetration, and peripheral side effects, have driven the development of various innovations aimed at enhancing its efficacy and tolerability. Among the most notable advancements are Temporal Interference (TI) stimulation and high-definition transcranial direct current stimulation (HD-tDCS), each of which employs unique principles to address these challenges.

Similarly to ISP, Temporal Interference (TI) stimulation, as proposed by Grossman and colleagues (Grossman et al., 2017), also utilizes the non-linearity of neuronal cell membranes (Luff et al., 2024). However, TI stimulates by the interference patterns of two concurrently applied high-frequency signals through separate electrode pairs. The interaction of these currents creates a low-frequency electric field at a targeted region, enabling low-frequency entrainment of deeper brain structures while exposing the superficial layers only to the high-frequency pattern. Clinical applications of TI have shown promise in treating conditions such as depression (Demchenko et al., 2024; Grossman et al., 2018) and have been proven to modulate brain activity in various models of health and disease (Beanato et al., 2024; Missey et al., 2021; Modak et al., 2024). However, TI relies heavily on precise electrode placement and stable interference conditions, which can be challenging in practical settings. Additionally, the high-frequency components can induce peripheral nerve stimulation, limiting tolerable intensities and constraining effective dose delivery to deeper targets.

High-definition tDCS, on the other hand, improves the spatial resolution of traditional tDCS by utilizing multi-electrode array configurations. This approach confines the electric field to smaller cortical areas, enhancing focality and reducing off-target effects (Datta et al., 2009). HD-tDCS has demonstrated its utility in modulating cortical excitability for cognitive enhancement and therapeutic applications, including stroke rehabilitation and depression (Bahr-Hosseini et al., 2023; Borckardt et al., 2012; Caparelli-Daquer et al., 2012; Donnell et al., 2015; Jog et al., 2023; Kuo et al., 2013; Nikolin et al., 2015; Shekhawat et al., 2015). Despite these advantages, HD-tDCS struggles with efficiently stimulating deeper brain structures.

In contrast, Intersectional Short-Pulse (ISP) stimulation uses a paradigm that combines temporal and spatial principles. By delivering rapid sequences of brief electric pulses through dynamically switching electrode pairs, ISP exploits the neuronal membrane’s capacity for temporal integration. This scalar integration mechanism enables ISP to achieve increased penetration to deep brain structures. Additionally, ISP does not introduce any constraint on the stimulation waveform envelope, allowing for precise temporal targeting of complex neuronal oscillations. The temporal and spatial distribution of charge injection reduces peripheral discomfort, facilitating the delivery of higher stimulation intensities and improving overall efficacy, making it suitable for phase-matched closed-loop applications such as seizure termination.

To further advance ISP and make informed decisions regarding optimal stimulation patterns and electrode configurations, a validated modeling pipeline is necessary. Additionally, a mechanistic understanding of how ISP interacts with neurons is critical for refining its application and maximizing its therapeutic potential.

### Validation of ROAST for ISP Stimulation

As a validation step, we compared the predictions of the ROAST (Realistic volumetric-approach to simulate transcranial electric stimulation) finite element modeling algorithm against actual measurements in human cadavers during ISP stimulation. This validation was essential to ensure computational model reliability in our ISP high-frequency regime.

The strong correlation between the modeled and measured voltage gradients provides robust support for the ROAST in simulating ISP-induced electric fields. This high degree of concordance across various stimulation patterns and brain regions, with accurate predictions for larger induced electric field gradients.

Our model predictions were less precise for stimulations between closely spaced electrodes, likely due to local tissue shunting effects and the model’s sensitivity to exact tissue segmentation. This observation highlights the importance of electrode placement when designing ISP protocols and interpreting simulation results.

The consistent correlation coefficients observed for ISP pairs with swapped electrode polarities further underscore the reliability of the ROAST framework for modeling transient electric fields. Swapped polarities also reduce undesirable side effects, such as scalp burning (Vöröslakos et al., 2018). These findings suggest that the model accurately captures the fundamental physics of current flow in the brain, regardless of the specific electrode configuration.

### Non-linear neuronal readout of the instantaneous extracellular electrical fields

Contrary to the expectation that increasing extracellular electric field (EF) intensity would linearly enhance neuronal depolarization, in our modeling work, we observed an initial somatic hyperpolarization under weak EFs that transitioned to action potential initiation as the EF intensity increased. This poses the issue of how mild hyperpolarization can lead to spiking with elevated stimulation. Excitable neurons exhibit a non-linear response to EFs, where the effect of the EF on neuronal activity depends on the intensity of the field and neuronal morphology. At subthreshold EF intensities, passive polarization can cause somatic hyperpolarization and distal depolarization (e.g., distal axonal segments or tuft dendrites). This occurs because neuronal compartments respond differentially to EFs due to variations in geometry and ion channel distributions. According to the model, as the EF intensity increases, the depolarization at excitable regions, such as the Ranvier nodes of distal axonal compartments, can reach the threshold for action potential initiation, leading to antidromic spiking even if the soma remains hyperpolarized. This non-linear readout underscores the need to consider spatially heterogeneous effects of EFs on neurons. The timing and spatial distribution of EFs significantly influence neuronal excitability (Radman et al., 2007), highlighting that weak fields can modulate neuronal activity when they interact with intrinsic neuronal properties. This emphasizes the complex interplay between extracellular fields and neuronal structures, with implications for understanding neural processing and for developing neuromodulation therapies.

### Non-linear integration of ISP-induced extracellular fields by neurons

Our computational modeling and *in vivo* patch-clamp recordings provide converging evidence that neurons integrate ISP-induced electric fields in a scalar rather than vectorial manner. This finding departs from the traditional understanding of how neurons respond to externally applied electric fields. The scalar integration allows ISP to overcome the "mirror effect" that limits conventional TES, where simultaneous activation of surface electrodes induces opposing effects under cathode and anode placements (Creutzfeldt et al., 1962).

Several key observations from our study support the scalar integration hypothesis. Our computational models revealed that the neuronal response to ISP stimulation correlated strongly with the scalar sum of individual ISP fields rather than their vectorial sum. This finding was corroborated by *in vivo* patch-clamp recordings, which showed that the scalar sum of pulse intensities more precisely explains the integrated effect of ISP stimulation. Further, the intensity threshold for evoking action potentials was similar for superficial and deep brain structures during ISP stimulation, in contrast to the depth-dependent effects seen with conventional TES. Collectively, these findings suggest that ISP pulse integration effects occur primarily due to the capacitive properties of the cell membrane rather than the instantaneous summation of extracellular electric fields. This mechanism allows ISP to achieve more uniform stimulation at different brain depths, an advantage for targeting deep brain structures such as the hippocampus.

### Network effects and seizure termination by ISP stimulation

While our study primarily focused on single-neuron responses, the efficacy of ISP in seizure control demonstrated significant effects on neural networks and brain oscillations as well. ISP can generate multidirectional electric fields in target structures that may disrupt synchronization across neuronal populations. By influencing the timing and probability of neuronal firing in a spatially focused manner, ISP may reset the phase of ongoing oscillations and prevent the spread of epileptiform activity. This network-level impact provides a crucial link between the observed cellular effects and the macroscopic seizure control.

The enhanced efficacy of closed-loop ISP stimulation in terminating seizures may result from precise temporal alignment with ongoing seizure rhythms. This timing could enable destructive interference with the pathological oscillations (Berényi et al., 2012; Takeuchi et al., 2021). By delivering focused, multidirectional stimulation pulses at specific phases of the seizure rhythm, ISP may disrupt the spatial and temporal coherence necessary for seizure maintenance and propagation. We hypothesize that the scalar integration of ISP-induced fields by neurons results in a more robust and widespread disruption of hypersynchronous activity compared to conventional TES. The rapid cycling of ISP stimulation sites may limit adaptive changes in neural networks that could reduce standard TES stimulation efficacy over time.

Future studies employing multi-site recordings or functional neuroimaging during ISP stimulation could elucidate these network-level mechanisms, potentially revealing how ISP modulates functional connectivity and oscillatory dynamics in both healthy and pathological states.

### Improved spatial focusing

The spatial focusing effect of ISP stimulation, as demonstrated by our simulations and *in vivo* experiments, overcomes a major limitation of conventional TES. By generating electric fields with similar strength and direction in bilateral deep structures (e.g., hippocampi), ISP can achieve targeted stimulation that is difficult to achieve with standard TES. This improved spatial specificity is key for targeting specific brain networks or structures (e.g., TLE).

The marked reduction in seizure duration and severity achieved by closed-loop ISP stimulation in our rat TLE model underscores ISP’s therapeutic potential. The rapid seizure termination and reduced intensity of motor manifestations is a dramatic advance over existing non-invasive stimulation techniques. Moreover, optimizing pulse duration without losing efficacy suggests that ISP protocols can be fine-tuned to minimize side effects while maintaining therapeutic benefits.

### Clinical translation and potential clinical applications

While our findings in the rat TLE model are promising, translating these results to humans is challenging. The human brain is larger with complex folding, making focused stimulation in deep structures more difficult. However, the scalar integration in our study should also apply to human neurons, suggesting that ISP could maintain its advantages over conventional TES in human applications. Computational models of ISP in human head models, followed by carefully designed clinical trials are crucial next steps. Initial human studies might focus on individuals with refractory epilepsy who are candidates for invasive treatments, allowing for direct comparison of ISP with other interventions. Our closed-loop ISP paradigm aligns well with emerging trends in personalized medicine, potentially allowing for tailored stimulation parameters based on individual seizure patterns. Moving to human trials, we need to optimize ISP parameters for the human brain, assess its safety profile over extended use, and explore its potential in treating other neurological and psychiatric disorders involving aberrant neural synchronization. The promising results of ISP stimulation in controlling epileptic seizures open up several avenues for clinical application and further research. ISP could be a non-invasive alternative or adjunct to treat drug-resistant epilepsy, particularly for patients who are not candidates for surgical interventions or who have not benefited from existing surgical and neuromodulatory approaches. Beyond epilepsy, non-invasive ISP for focused, deep brain stimulation could help treat neuropsychiatric disorders involving limbic and extrapyramidal structures, including major depressive disorder, anxiety disorders including obsessive-compulsive disorder, and Parkinson’s disease. Future studies should explore the potential for patient-specific ISP protocols, optimizing electrode placement and stimulation parameters based on individual brain anatomy and pathology. Integrating ISP with individual structural and functional neuroimaging could further enhance the targeting for spatial and temporal precision. While our study focused on acute effects, future research should investigate the long-term impacts of ISP stimulation on brain plasticity and potential beneficial as well as adverse effects.

### Limitations

Despite the promising results, several limitations of our study require further research. While the rat kindling model is well-established, studies in larger animals and eventually humans are necessary to confirm or refute the translational potential of ISP. Although we propose scalar integration as the primary mechanism, further investigation into the cellular and network-level effects of ISP is warranted. While we explored pulse duration, other parameters, such as frequency, intensity, and electrode configuration, could be further optimized. Finally, a direct comparison of ISP with other emerging TES approaches, such as temporal interference stimulation, is needed to fully assess its relative advantages and potential clinical utility.

In conclusion, ISP stimulation is a significant advancement in non-invasive brain stimulation. By exploiting the scalar integration of induced electric fields by neurons, ISP improves spatial focusing and efficacy in modulating neural activity, particularly in deep brain structures. Our ability to rapidly terminate temporal lobe seizures in a rat model highlights its compelling therapeutic potential. With additional refinements and validation in humans, ISP stimulation may open new avenues to treat diverse neurological and psychiatric disorders, offering hope for patients who have not responded to conventional therapies.

## Acknowledgments

We are grateful for Márk Harangozó and András Kispál for their technical assistance and György Buzsáki and Mihály Vöröslakos for their constructive discussions on the manuscript. This work was supported by the Momentum program II of the Hungarian Academy of Sciences (AB), EFOP-3.6.1-16-2016-00008 (AB), EFOP 3.6.6-VEKOP-16-2017-00009 (AB), and KKP133871/KKP20 grants of the National Research, Development and Innovation Office, Hungary (AB), the 20391-3/2018/FEKUSTRAT of the Ministry of Human Capacities, Hungary, the EU Horizon 2020 Research and Innovation Program (No. 739593— HCEMM to AB), Ministry of Innovation and Technology of Hungary grant (TKP2021-EGA-28 to AB), Hungarian Scientific Research Fund (Grants NN125601 and FK123831 to MLL, K135837 to ZS), the Hungarian Brain Research Program (grant NAP2022-I-7/2022 to AB & MLL), UNKP-20-5 New National Excellence Program of the Ministry for Innovation and Technology from the source of the National Research, Development and Innovation Fund (MLL), Hungarian Research Network TECH-2024-020 to ZS.

## Author Contributions

A.B. conceived the project.

T.F., M.S., Z.Ch., B.R., B.H., E.V., I.L., P.R., A.P., L.B., G.K., N.F., O.N., T.L., M.L.L. and A.B. performed the experiments.

T.F., M.S., A.J.N. and A.B. analyzed the data.

T.F., K.F., Z.S. and A.B. performed the simulations.

T.F., M.S., M.L.L., O.D., and A.B. wrote the manuscript.

M.L.L. and A.B. supervised the project.

## Materials and methods

### Model neurons and E-field stimulation of model neurons

We adjusted models of neurons with realistic axonal and dendritic morphologies for simulations of the single-cell response to stimulation using the electric field #15 (basket) and #20 (pyramidal) cells (Aberra et al., 2018). The field intensity (ISP intensity) was scaled to determine the threshold to evoke one AP with a whole sequence of ISP pulses of six different pairs of sources. We used NEURON software (Carnevale & Hines, 2006) and customized Python (3.8.8 on macOS Sonoma) scripts to model the effect of local extracellular fields with gradually changing intensities and directions (representing the local effects of the consecutive ISP pulses).

### Simulating transcranial electric stimulation (ROAST)

The finite element model (FEM) model for electric field simulation was created from rat head MR scans with ROAST modeling software. Models are constructed based on magnetic resonance images (MRI) of the rat head (Goerzen et al., 2020) for a detailed representation of the three-dimensional anatomy. To simulate current flow on an MRI is segmented, virtual electrodes are located on this anatomical model, the volume is tessellated into a mesh, and the FEM is solved numerically to estimate the current flow. We used an open-source implementation called the Realistic volumetric-approach to simulate transcranial electric stimulation (ROAST). ROAST combines the segmentation algorithm of SPM12, a Matlab script for touch-up and automatic electrode placement, the finite element mesher iso2mesh and the solver getDP (Huang et al., 2019).

### Cadaver recordings

The cadaver experiments were approved by the Regional and Institutional Review Board of Human lnvestigations in University of Szeged (22/2023-SZTE) and the Medical Research Council - Scientific and Research Ethics Committee of Hungary (BM/18167-3/2023). Two cadavers were investigated in this study. Cadavers were selected based on the absence of significant skull or brain abnormalities and recent anatomical imaging (CT or MRI) to ensure accurate modeling of electric current flow. After positioning the cadaver prone on the autopsy table, standard skull measurements were recorded, and a 10-20 EEG electrode cap was used to mark the electrode entry points. Four electrode strips (Ad-tech TS08R-SP10X-000) were implanted subgaleally through small incisions, secured with sutures, and verified by palpation. A stereotactic frame was attached symmetrically to the skull using ear bars, and the stereo EEG (sEEG; Medtronic 3389-28) lead insertion was guided by predefined trajectories planned via merged cadaver imaging with the stereotactic frame model. The sEEG leads were stained with ink to mark the electrode tracks for post-experimental track reconstruction, ensuring accurate analysis of their trajectories in subsequent histological evaluations. Positions of the sEEG leads and the subgaleal strips were verified by a C-arm X-ray. Electrical recordings were obtained at multiple depths.

ISP stimulation was delivered through the subgaleal strip electrodes using the Neunos Neuroclinical device (Neunos, Szeged, Hungary), while biosignal recordings were performed from the sEEG electrodes with the Amplipex KJE-1001 data acquisition system (Amplipex, Szeged, Hungary), with a sampling rate of 640 kS/s per channel and no signal filtering. Impedance checks were performed at each depth, and predefined ISP stimuli were delivered at the strip electrodes.

After the recordings, the brain was removed and fixed in a paraformaldehyde bath for at least two weeks before sectioning. Macroscopic analysis of the sEEG lead trajectories was conducted on coronal slices of the fixed brain. The electrode and lead positions were reconstructed using pre-mortem imaging data and Slicer3D software.

### Animals

All experiments were performed following European Union guidelines (2003/65/CE) and the National Institutes of Health Guidelines for the Care and Use of Animals for Experimental Procedures. The Ethical Committee approved the experimental protocols for Animal Research at the Albert Szent-Györgyi Medical and Pharmaceutical Center of the University of Szeged (XIV/824/2021). Ten male Long-Evans rats were used in the experiments.

### Surgery

The animals were operated on under isoflurane anesthesia. The rats were implanted with recording electrode triplets (inter-wire spacing, 0.4 mm) (Kozák et al., 2018) the frontal areas (AP: +2 mm from the bregma; ML: +2 mm in the sagittal plane; DV: -1,4 mm from the dura) and the dorsal hippocampus (AP: -4 mm from the bregma; ML:+0.7 mm, +2.7 mm, +4 mm in the sagittal plane; DV: -3 mm from the dura) bilaterally. Five pair stimulation stainless steel screw electrodes were implanted over the temporal bone bilaterally. Miniature stainless-steel screws (serving as reference and ground) were implanted above the cerebellum. A copper mesh (acting as a Faraday cage) was built around the electrode holders and enforced with dental cement (Kozák et al., 2018). A custom-built bipolar electrode consisting of two insulated tungsten wires was placed targeting the hippocampal commissure for kindling and seizure induction during the experiments.

### Hippocampal electrical kindling

A bipolar stimulus electrode for kindling and seizure induction was prepared as described above. The bipolar stimulus electrodes’ (inter-wire spacing, 0.5 mm) insulations 0.4-0.5 mm around the tips were stripped to decrease their impedance to 10-20 kΩ at 1kHz.The stimulus electrode was implanted on the HPC commissure (AP: -0.84 mm from the bregma; ML: 1.5 mm, angled at 18° from the sagittal plane; DV: -1.5 mm). After recovery from the implantation surgery, the HPC was stimulated every day at subconvulsion intensity via the kindling electrode. Each kindling stimulation included 120 × 0.5 ms positive – 0.5 ms negative bipolar rectangular pulses at 62.5 Hz (Gelinas et al., 2016) delivered by an isolated stimulator generator in voltage control mode (STG4008; Multi-Channel Systems). Stimulus intensity was set as the minimum induced after-discharge (10-25 Hz population spikes synchronously recorded in > 50% of HPC channels after the HPC commissure stimulation), commonly ±600-3000 mV. The kindling stimulation was performed six times per day for ten days. The interstimulus intervals were at least 30-min.

### Data acquisition from freely moving rats

Local field potential (LFP) recordings were made in the animals’ home cages, and food and water were given *ad libitum*. The recording wire electrodes were connected to a signal multiplexing headstage (HS3_v1.3, Ampliplex) and stored after digitalization at a 500 Hz sampling rate (KJE-1001, Ampliplex) (Berényi et al., 2014). In parallel, rats’ preamplified signals were analyzed online by a programmable digital signal processor (RX-8, Tucker-Davis Technologies, FL, USA) using a custom-made seizure detection algorithm (Kozák & Berényi, 2017).

### Closed-loop transcranial stimulation

HPC electrographic seizure waves were detected online using a programmable digital signal processor. These detected waves triggered the transcranial stimulation for on-demand real-time seizure interventions (Kozák & Berényi, 2017). The LFP signals were analyzed online to detect LFP deflections in the HPC using a custom-made seizure detection algorithm based on a previously developed method. Briefly, the pre-selected HPC channel was band-pass filtered with a 4^th^-order Butterworth filter, rectified, and integrated into a time window. Each transcranial stimulus was triggered by an ictal HPC LFP deflection when the filtered signal exceeded a threshold. The time resolution of the detection was 2 ms. Synchronous multiple threshold crossings triggered a trapezoidal single-pulse (0.5 ms; 1 ms; 5 ms; 15 ms; 80 ms) stimulation (STG4008; Multi-Channel Systems). The stimulation was performed in voltage mode. In conventional TES or ISP mode (see ‘Introduction’ section), the stimulus intensity was 8000 mV. For ISP stimulation, each electrode pair was pulsed for 2.5 μs, and the pulses cycled through the three pairs for 500 ms followed by non-stimulated 1-s control periods. This sequence was repeated to stimulate the right or left hemisphere alternatingly. Diffuse stimulation was applied between the left and right temporal electrodes. On each experimental day for the closed-loop intervention trials, we performed at most six recorded induced seizures. At most, two sessions of each type (sham, ISP, diffuse) were conducted. Some days, we could only perform less than five recordings. The order of types was randomly chosen daily. For the ISP pulse duration parameter optimization experiment, we tested every predetermined parameter daily in random order. To measure the electric field, 100 stimulation cycles were averaged for both conditions and then the peak-to-peak amplitude was measured for each channel. The first spatial derivate of stimulus-induced voltage was calculated.

### Behavioral monitoring

The rats’ behavior was continuously monitored during the electrophysiological recordings with a webcam (LifeCam Studio) and synchronized with the recorded neuronal data. The severity of motor seizures was estimated online and offline based on Racine’s scale for seizures: 1, mouth and facial movements; 2, head nodding; 3, forelimb clonus; 4, rearing; 5, rearing and falling (Racine, 1972).

### Histology

After data acquisition, electrolytic lesions were made at the electrodes’ tip to verify their location and possible pathologic changes. Rats were deeply anesthetized with 1.5 g/kg urethane (intraperitoneal) and transcardially perfused with saline, followed by 4% paraformaldehyde (PFA) and 0.2% picric acid (PA) in 0.1 M phosphate buffer. Brains were post-fixed overnight, and 50 μm-thick coronal sections were prepared with a vibratome (VT100S Vibratome, Leica) and stained with 4’,6-diamino-2-phenylindole dihydrochloride (DAPI; D8417, Sigma-Aldrich).

### *In vivo* whole-cell patch-clamp recordings with concurrent transcranial electrical stimulation

A 3-month-old female Wistar rat was anesthetized with urethane and was implanted with recording electrode triplets as described above. A craniotomy of 1 mm in diameter was drilled above the somatosensory cortex of the right hemisphere to provide access to the brain while leaving the dura intact. A small sink was built from dental cement surrounding the craniotomy and was filled with ACSF to prevent the tissue from drying and holding the ground electrode in place. Pipettes were pulled from thick-walled borosilicate glass (BF150-86-15, Sutter Instrument, Novato, CA) with a Sutter P-1000 puller (Sutter Instrument, Novato, CA). Pipette tip resistance ranged between 5-10 MΩ when the electrodes were filled with a K-gluconate-based internal solution (containing, in mM: 130 K–gluconate, 8 KCl, 10 HEPES, 4NaCl,4Mg-ATP, and 0.3Tris-GTP,14(tris-)phosphocreatine) with pH = 7.28 and osmolality 295 mOsm) at 32°C). Whole-cell patch-clamp recordings were performed blindly 200-250 μm under the dura, most likely targeting L2/3 pyramidal neurons. Recordings were discarded if access resistance exceeded 60 MΩ or if the holding current at -60 mV was larger than -500 pA. Signals were amplified using a MultiClamp 700B amplifier and were digitized at a sampling frequency of 50 kHz using an Axon Digidata1440 digitizer that was controlled by pClamp 10.3 software (Molecular Devices, San Jose, CA). ISP stimulation epochs were manually delivered while in whole-cell configuration in current clamp mode at an intensity of 100 μA and a duration of 50 ms (17-18 times). The same ISP stimulation sequences were repeated extracellularly as well (20 times) to compare the induced electrical artifacts between the two configurations. Recordings were subsequently processed with the pyABF Python site-package (Harden, 2022) and analyzed offline by custom-written Python scripts.

### Analysis of cadaver recordings and corresponding simulations

To validate the *in silico* ROAST modeling of rapid transient fields induced by ISP stimulation, we compared the simulated voltage gradients with real-life measurements recorded from cadavers. The ROAST model incorporated MRI-based electrode positions and the precise ISP stimulation sequence delivered via subgaleal electrode strips, identical to those used in the cadaver measurements. This allowed for the generation of 3D potential maps, which provided simulated voltage values at predefined voxel locations corresponding to the sEEG electrode contact sites.

The sEEG electrode recordings obtained from cadaver experiments were processed to extract discrete voltage potentials for each 100-µs ISP slot. Post-processing of the recorded data involved detrending, aligning, and extracting the median values across multiple repetitions of the ISP stimulation. These extracted values were used to construct potential maps, which represented the measured voltage gradients at the sEEG electrode contact points.

The next step in validation involved mapping the measured electrode positions from the cadaver recordings to the corresponding voxel locations in the ROAST model. Pairwise voltage differences were calculated for each electrode pair, generating a voltage difference matrix for both the simulated and real-life data. To assess the accuracy of the ROAST model, we performed pairwise correlations between the simulated and recorded voltage gradients. A matrix of correlation values was created for all possible sEEG electrode pairs, allowing for a detailed comparison between the model and experimental results.

A more granular analysis was also performed to investigate the correlation between the simulated and measured values as a function of the strength of the voltage gradient and as a function of the distance between the active anodes and cathodes. It was hypothesized that smaller voltage gradients would exhibit a lower SNR, potentially diminishing the accuracy of the predictions. Additionally, it was assumed that more localized stimulation would generate strong electric fields in restricted brain regions, which could result in many gradients being near zero or noisy.

### Duration of electrographic seizures and normalized seizure length

Signals sampled at 500 Hz were filtered between 1 and 250 Hz for LFP signals. Peri-stimulus LFPs (30 s baseline and 180 s test epochs) were then extracted using timestamps recorded in a digital channel. The peri-stimulus LFP signals were further band-pass filtered to 10–80 Hz to prepare narrow-band LFPs. The narrow-band LFPs were then smoothed using a three-second-long moving average filter. The duration of the HPC and CTX electrographic seizures were automatically detected and defined in test epochs when all amplitudes of the smoothed LFPs in each brain region exceeded three times the root-mean-square levels of the corresponding baseline epochs. Electrographic seizures’ durations with electrical interventions were refined by the consensus estimate of two experienced researchers doing manual inspections and using Neuroscope software (RRID: SCR_002455) (Hazan et al., 2006). This refinement was done because the automated detection algorithm sometimes misestimated durations due to electrical artifacts. The variation between the manual estimates of both researchers and the automated detection data of ISP interventions was less than five percent. We normalized the seizure length to each day’s most prolonged seizure duration.

### Assessment of the severity of motor seizures

The severity of motor seizures was video monitored and evaluated according to Racine’s scale on each trial.

### Statistical analysis

All data analyses were performed in MATLAB (RRID: SCR_001622; Mathworks, Natick, MA, USA) unless otherwise noted. Values are expressed as mean ± standard deviation (SD) unless otherwise stated. MATLAB’s Statistics Toolbox (Berens, 2009) was used for the statistical tests. Effects of the treatment on the seizure duration and Racine’s score values were tested using Wilcoxon signed-rank test. Friedman test followed by Nemenyi test was used to test the effect of ISP pulse duration. The significance level was set at *p*<0.05. One, two and three asterisks indicate significance levels <0.05, <0.01, and <0.001, respectively.

## Data Availability

The data generated in this study are available from the corresponding author upon reasonable request.

## Code Availability

All custom codes are freely available from the corresponding author on reasonable request.

