## Supplementary Figure for "Non-vectorial Integration of Intersectional Short-Pulse Stimulation Enables Enhanced Deep Brain Modulation and Effective Seizure Control"

[9]Department of Neurology, NYU Langone Comprehensive Epilepsy Center, NYU Grossman School of Medicine, New York, NY 10016, USA

**Supplementary Figure 1.**

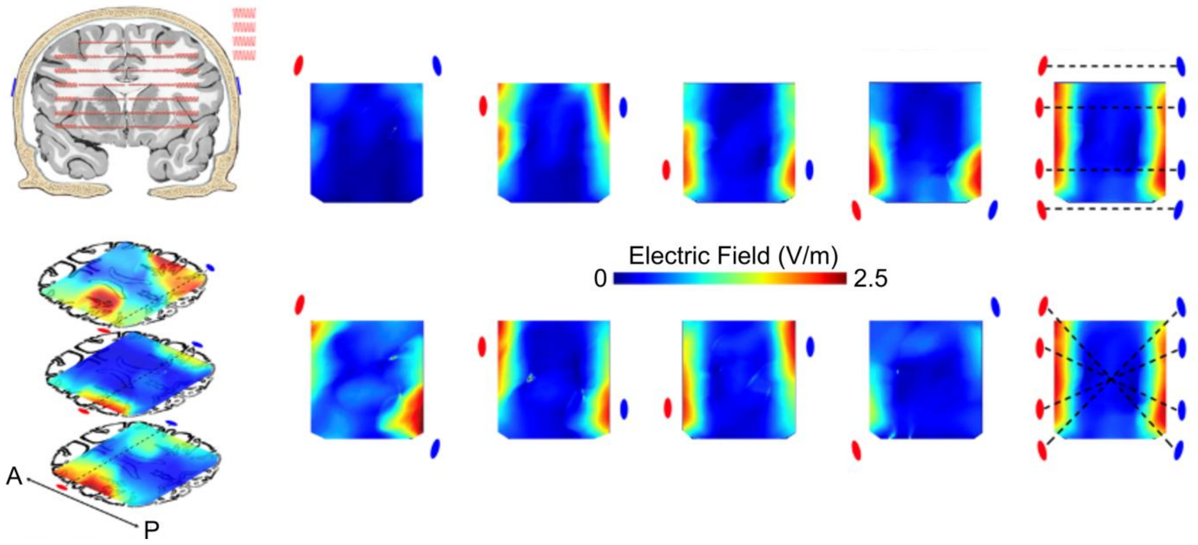


**Supplementary Figure 1: Equivalence of intracerebral electric field distributions in human cadaver measurements under different transcranial stimulation configurations.**

This figure demonstrates the experimental validation of the principle that different electrode configurations produce identical electric field distributions in the brain when applied simultaneously. The experiment was conducted using human cadaver specimens. For detailed methodology, see (Vöröslakos et al., 2018). Left: Schematic representation of the experimental setup. Top image shows a coronal section of the human head with implanted sEEG electrodes for intracerebral field measurements. Below are three axial slices showing the distribution of the intracerebral electric fields, recorded by the electrodes. Right: Measured electric field distributions for five different stimulation configurations. Each column represents a different electrode arrangement, with red and blue circles indicating the positions and polarity of the transcranial electrodes. The rightmost column shows the cases when all independent sources are applied simultaneously. Top row: sources are connected in a non-crossing (i.e. linear) fashion. Bottom row: Electrodes are assigned to the sources in a way to cross each other. Despite the different electrode configurations (crossing vs. linear arrangements of the four independent current sources), the resulting intracerebral electric field distributions are nearly identical when all sources are activated simultaneously. This is evidenced by the comparable patterns and magnitudes of electric fields across stimulation conditions.

**Supplementary Figure 2.**

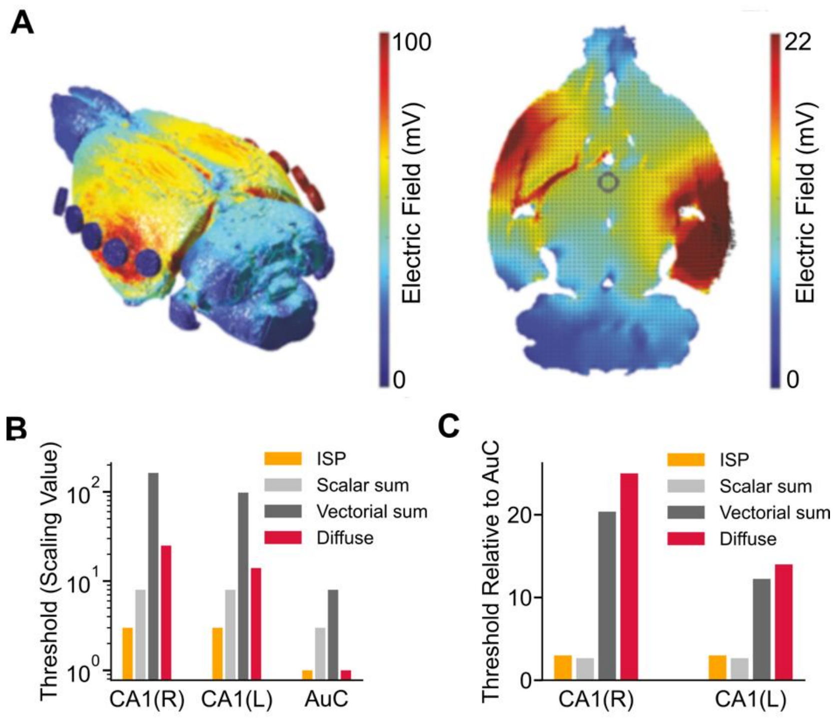


**Supplementary Figure 2: Simulation of ISP and diffuse TES effects on single neurons in the rat brain. (A)** Model of rat brain, with 1mA current injected at one electrode pair (i.e. one timeslot of the ISP stimulation sequence; left: 3D model, right: transverse plane). **(B)** The stimulus intensity threshold for ISP entrainment (orange) is similar near the electrodes (auditory cortex, AuC) and at the targeted deep crossing points (left and right hippocampal CA1). In contrast, conventional TES (red) requires substantially higher intensity to entrain deep structures than at sites proximal to the stimulating electrodes. The integrated ISP effect can be approximated better with the scalar sum of individual ISP pulses (light grey) than with the vectorial sum (dark grey). **(C)** Same as (B), relative to surface-proximal structures (AuC).

**Supplementary Figure 3.**

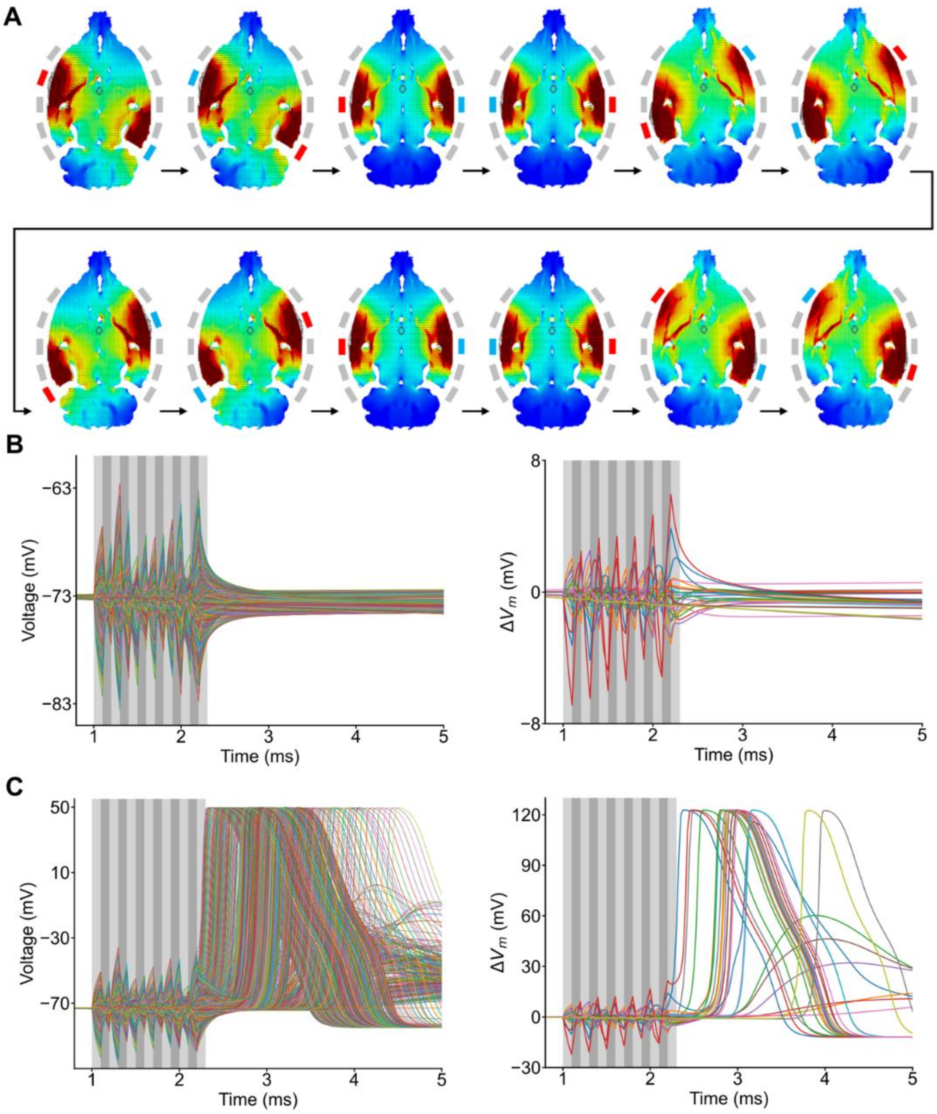


**Supplementary Figure 3: Simulation of ISP and diffuse transcranial electrical stimulation effects on single neurons in the rat brain. (A)** Induced, instantaneous electric fields in a rat brain, with current injected at one electrode pair at a time (i.e. one timeslot of the ISP stimulation sequence). Blue and red electrodes around the brain sections denote the active anode and cathode in each case, while the inactive electrodes are shown in gray. **(B** and **C)** Simulation of the effect of the same symmetric ISP stimulation sequence on a model neuron demonstrates the same non-linear behavior, where the effects of mirrored electric fields do not cancel out. Panels on B and C denote subthreshold and suprathreshold stimulations, respectively. While the left panels display the potential traces of every modeled segment of the realistic neuron model, the right panels show only a zoomed subset of them for better visibility.
